# Changes in blood cell deformability in Chorea-Acanthocytosis and effects of treatment with dasatinib or lithium

**DOI:** 10.1101/2021.12.16.472921

**Authors:** Felix Reichel, Martin Kräter, Kevin Peikert, Hannes Glaß, Philipp Rosendahl, Maik Herbig, Alejandro Rivera Prieto, Alexander Kihm, Giel Bosman, Lars Kaestner, Andreas Hermann, Jochen Guck

**Affiliations:** Max-Planck-Institut für die Physik des Lichts and Max-Planck-Zentrum für Physik und Medizin, Erlangen, Germany; Biotechnology Center, Center for Molecular and Cellular Bioengineering, Technische Universität Dresden, Dresden, Germany; Translational Neurodegeneration Section “Albrecht Kossel”, Department of Neurology, University Medical Center Rostock, University of Rostock, Rostock, Germany; Division for Neurodegenerative Diseases, Department of Neurology, Technische Universität Dresden, Dresden, Germany; Department of Experimental Physics, Saarland University, Campus E2 6, Saarbrücken, Germany; Department of Biochemistry, Radboud UMC, Nijmegen, The Netherlands; Theoretical Medicine and Biosciences, Saarland University, Building 61.4, 66421 Homburg, Germany; Deutsches Zentrum für Neurodegenerative Erkrankungen (DZNE), Rostock/Greifswald, Rostock, Germany; Center for Regenerative Therapies Dresden, Technische Universität Dresden, Dresden, Germany

**Keywords:** chorea-acanthocytosis, blood cell deformability, real-time deformability cytometry, dasatinib, lithium, cell mechanics

## Abstract

Misshaped red blood cells (RBCs), characterized by thorn-like protrusions known as acanthocytes, are a key diagnostic feature in Chorea-Acanthocytosis (ChAc), a rare neurodegenerative disorder. The altered RBC morphology likely influences their biomechanical properties which are crucial for the cells to pass the microvasculature. Here, we investigated blood cell deformability of 5 ChAc patients compared to healthy controls during up to one-year individual off-label treatment with the tyrosine kinases inhibitor dasatinib or several weeks with lithium. Measurements with two microfluidic techniques allowed us to assess RBC deformability under different shear stresses. Furthermore, we characterized leukocyte stiffness at high shear stresses. The results show that blood cell deformability – including both RBCs and leukocytes - in general is altered in ChAc patients compared to healthy donors. Therefore, this study shows for the first time an impairment of leukocyte properties in ChAc. During treatment with dasatinib or lithium, we observe alterations in RBC deformability and a stiffness increase for leukocytes. The hematological phenotype of ChAc patients hints at a reorganization of the cytoskeleton in blood cells which partly explains the altered mechanical properties observed here. These findings highlight the need for a systematic assessment of the contribution of impaired blood cell mechanics to the clinical manifestation of ChAc.

## 1 Introduction

Chorea-acanthocytosis/VPS13A disease (ChAc) is a rare monogenetic neurodegenerative disease of the young adulthood affecting multiple systems other than the central nervous system, e.g., the red blood cells (RBCs) (1–3). ChAc is characterized by various movement disorders (due to a degeneration mainly of striatal neurons), epilepsy, cognitive decline and misshaped spiky RBCs, the latter being referred to as acanthocytes (1,2,4,5). The autosomal-recessive condition is caused by loss of function mutations in the *vacuolar protein sorting 13 homolog A* (*VPS13A*) gene (6–10). Therefore, the term “VPS13A disease” has been recommended to replace the historical, more descriptive terminology (11). A disease-modifying therapy has not been established so far.

It is still unclear, how exactly the loss of function of VPS13A leads to the manifestation of the cellular and clinical phenotype. VPS13A localizes at different membrane contact sites and is most probably involved in non-vesicular lipid transport as it is assumed also for the other members of the VPS13 protein family (for review, see Leonzino, Reinisch and De Camilli, 2021) (12–15). Two downstream mechanisms are considered to be important drivers of ChAc pathophysiology: decreased phosphoinositide-3-kinase (PI3K) signaling and increased activity of Src family tyrosine kinase Lyn (for review, see Peikert et al. 2018 (2)).

Active Lyn kinase accumulates in ChAc RBCs and hyperphosphorylates membrane proteins such as band 3 protein. Since band 3 is a structural protein, linking the cytoskeleton to the plasma membrane, it is likely to be causally involved in the genesis of the spiky morphology. Accumulation of active Lyn kinase was also found to be related to impaired autophagic flux (16,17). These phenotypes could be reversed by *in vitro* treatment with tyrosine kinase inhibitors (TKI) such as the Src kinase family inhibitor dasatinib (16). In a Vps13a knockout mouse model, the more specific Lyn kinase inhibitor nilotinib improved both the hematological and neurological phenotypes by improving autophagy and preventing neuroinflammation (18). In their recent work, Peikert et al., 2021 (19) report on single individual treatment approaches targeting Lyn kinase with dasatinib in ChAc patients. They showed that initially reduced F-actin signal, increased osmotic fragility and impaired autophagy were partially restored in ChAc RBCs.

Furthermore, disturbed signaling via Phosphoinositide-3-kinase (PI3K) has been also identified as relevant in the pathophysiology of ChAc patients. Reduced signaling via the PI3K-Rac1-PAK pathway was reported to lead to disordered actin polymerization (20). Additionally, in other cells, altered PI3K signaling caused a decreased ORAI1 expression and store-operated Ca^2+^ entry (SOCE), subsequently disturbed Ca^2+^ homeostasis and apoptosis which could be partly reversed by *in vitro* lithium treatment (21–23).

The rather late onset of the disease at the age of early adulthood and the slow progression rate (1,3,24) lead to the assumption of a dependency between the altered RBC properties and neuronal degeneration: Changes in the mechanical properties of the RBCs and hence reduced passage in micro-vessels and capillaries may impair oxygen supply in certain areas of the brain that could over time accumulate to tissue alterations leading to the symptoms described (25).

Altered RBC morphology is often linked to a change in cell mechanics which may lead to impaired blood flow (26,27). Until now, the effect of acanthocytes in ChAc patients on the pathophysiology of the disease remains elusive. Here, we describe, for the first time, altered deformability of blood cells from ChAc patients including white blood cells (WBCs). In a second step, we observed these parameters during individual off-label treatments with dasatinib or lithium, each targeting one of the signaling pathways believed to be involved in the genesis of the RBC phenotype.

Deformability was measured using two techniques that asses the response to different ranges of shear stresses acting on the cells: shape analysis of RBCs at low shear stress (0.1-3 Pa) (28) and real-time fluorescence and deformability cytometry (RT-FDC) (29,30) at high shear stress (ca. 100 Pa). RBC shape analysis only assesses the deformability of RBCs because the stresses are too low to deform WBCs and the throughput is too low to measure a sufficient number of WBCs. In contrast, RT-FDC was used to characterize the mechanics of both, RBCs and WBCs. In RT-FDC on RBCs, the interpretation of the deformation data is not as straight forward as for the RBC shape analysis because the RBCs undergo a non-trivial shape change here. We chose to use both methods to have a comprehensive image of the blood cell deformability at different stresses. We demonstrate that the deformability of all blood cells is affected in ChAc patients and this is further modulated by the treatments. This highlights that monitoring blood cell mechanical properties of ChAc patients during the course of disease and possible treatments can increase our understanding of this disease.

## 2 Materials and Methods

### 2.1 Cell source and reagents

We included 5 ChAc patients in this study for whom diagnosis was confirmed by Western blot (absence of chorein/VPS13A protein) and genetic testing (31). The clinical parameters of the patients are listed in Table S1 and complete blood count, RBC indices and hemolytic parameters are listed in Table S2. Further information for P1-P3 can be found in Peikert et al., 2021 (19). Patients and healthy control blood donors were enrolled in ongoing studies on the pathogenesis and natural history of neurodegenerative diseases approved by the institutional review board of the Technische Universität Dresden, Germany (EK 45022009, EK 78022015). The ChAc patients were treated with dasatinib (P1-P3) or lithium (P4-P5) in the context of an individual off-label therapy based on the above described preclinical evidence. Standard dose of 100 mg dasatinib per day was administered orally, as it was lithium for which serum lithium concentration target range was defined as 0.6-0.8 mmol/l. Treatment started after the blood for the baseline measurements was taken.

### 2.2 RBC shape analysis at low shear stress

The distribution of shapes for RBCs flowing through a narrow capillary at a fixed flow condition is directly linked to the distribution of shear moduli within the population. Thus, changes in the shape distribution are directly linked to a change of cell mechanics in the sample. This approach and the experimental design are described in detail in Reichel et al., 2019 (28). Here, cells flow through a roughly 5 mm long measurement channel that is 10 µm wide, and their shapes and dynamics are tracked at the end of the channel over a length of 435 µm. Cells were recorded with the same setup used for real-time deformability cytometry (29) using a different microfluidic chip. Examples of cells flowing through the region of interest are shown in figure 1A,B and in supplementary videos S1-S8.

**Figure 1:**
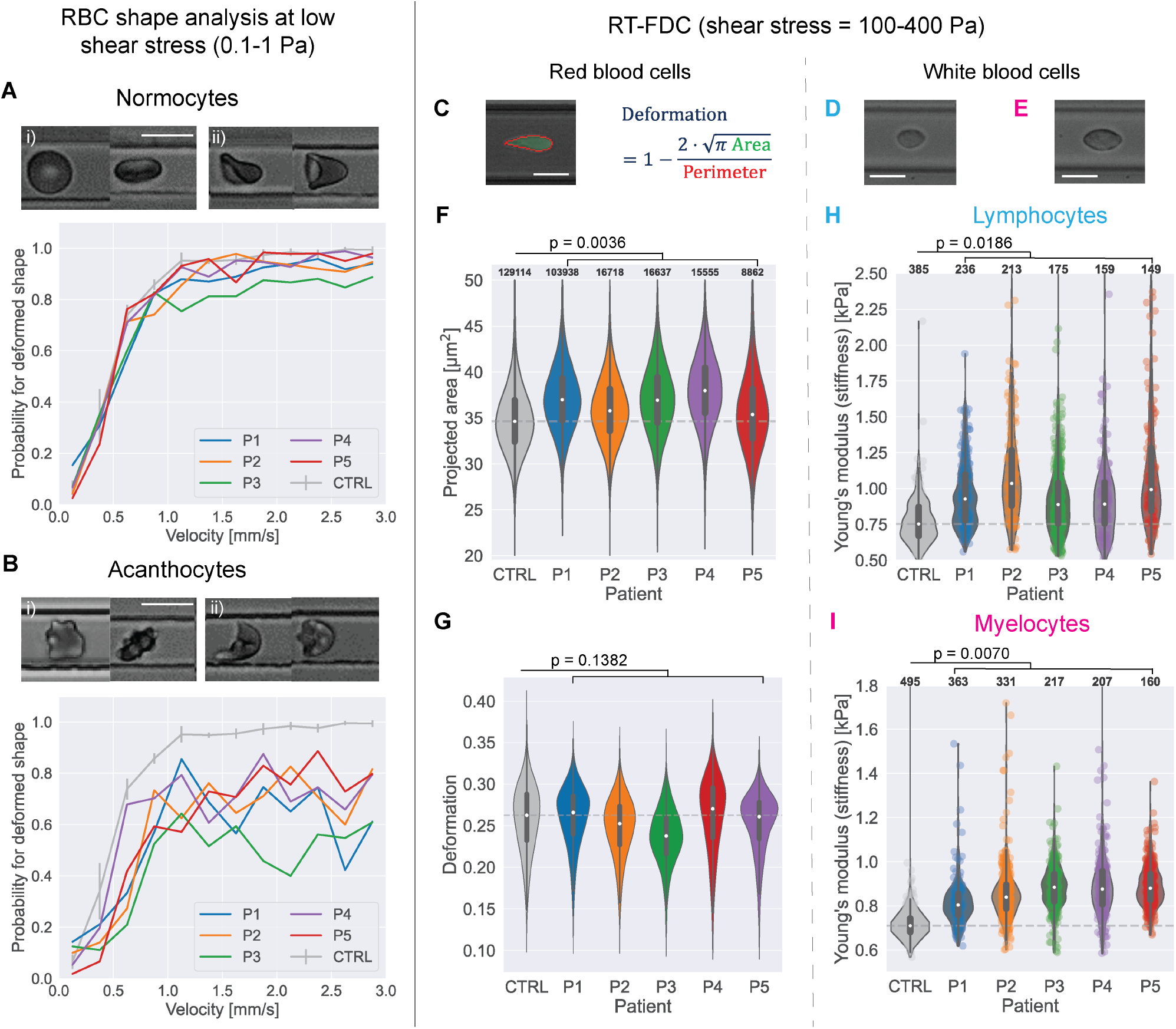
Blood cell deformability of ChAc patients compared to healthy control blood. **(A)** Example shapes of normal, discocyte RBCs (normocytes) in an **(i)** undeformed state and **(ii)** deformed by flow during low shear stress flow in a 10 µm channel. The diagram shows the probability to find normocyte cells from the sample in a deformed state at a given cell velocity in the channel for the acanthocyte patients and the mean curve from 3 control measurements. Error bars represent SEM. scale bar represents 10 µm **(B)** Example shapes of acanthocytes in an **(i)** undeformed state and **(ii)** deformed by flow. Below is the shape probability diagram for the acanthocytes and the control curves. **(C)** Example shape of an RBC in RT-FDC and representation how the deformation parameter is computed. All scale bars represent 10 µm. **(D)** Example image of a lymphocyte in RT-FDC. **(E)** Example image of a myelocyte. **(F), (G)** Projected area within the contour and deformation of RBCs of the ChAc patients from RT-FDC measurements vs. pooled data from 10 control measurements (full data in fig. S4). White dots represent the median value; grey box in the violin shows inter-quartile range (IQR) and extended lines 1.5×IQR. Dashed grey line shows the median control value. P-values were calculated with linear-mixed effect models as described in Herbig et al., 2018 (36). Numbers on top of the plots indicate the number of observations per violin. **(H)** Young’s modulus of lymphocytes from ChAc patients vs. control (n=1). **(I)** Young’s modulus of myelocytes from ChAc patients vs. control (n=1).

The shape analysis introduced in Reichel et al., 2019 (28), comprising the RBC shapes tumblers, tank-treaders, parachutes and multilobes, cannot directly be used for samples containing acanthocytes because their shape at rest differs from that of healthy RBCs. Further, acanthocytes show shapes not observed for healthy cells (discocytes or normocytes) when deformed by the flow in the channel. For this study, cells flowing through a square channel with a width of 10 µm at cell velocities ranging up to 3 mm/s were characterized by their membrane morphology as normal looking (referred to as normocytes) or as acanthocytes if they showed thorn-like protrusions. We did not further distinguish between acanthocytes and echinocytes but argue that the mechanical properties of both cell types should be very close (32). Furthermore, cells were classified as either deformed by the flow or still maintaining their resting shape (undeformed). This classification was done by eye. We chose the channel dimension of 10×10 µm because in such channels, RBCs undergo a sharp transition from undeformed discocyte to parachute shape with a low probability of other transient shapes (28) which makes it well suited to describe the deformed state of the cells. A change of the cell mechanical properties should manifest in a change of the fraction of deformed cells in the sample and that the velocity at which the majority of RBCs get deformed is shifted to higher values. For examples see figure 1A,B.

The classification was done by eye from the recorded videos of cells passing the region of interest. This resulted in a curve with probabilities to find cells in a deformed state for a given range of velocities. The resulting curve for healthy RBCs is given in figure 2A (data from Reichel et al., 2019 (28), 3 control samples). To characterize differences to ChAc patient cells and changes on the cells during treatment, a limited exponential growth function *P* = 1 − *e*^− *λ·υ*^, with cell velocity υ, was fitted to the data. With this function, the probability to deform at υ = 0 is zero and for very high velocities the probability converges to 100% which motivated this function because cells will eventually deform at high velocities. The cell deformability is characterized with the *growth rate λ* which describes how quickly the cells reach a deformed state. The fitted curve for the control values is also depicted in figure 2A. A higher growth rate would indicate that cells are more deformable.

**Figure 2:**
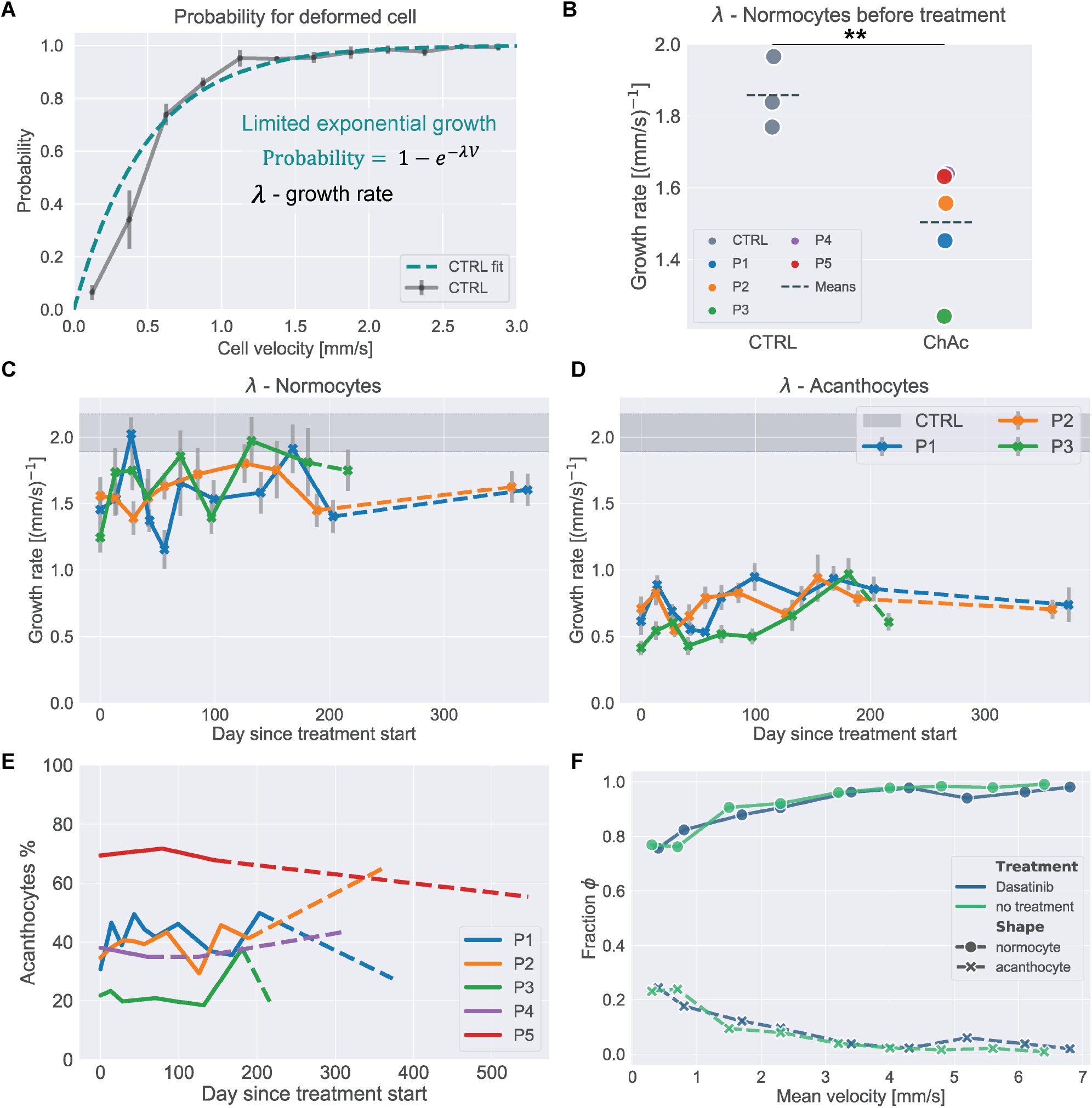
Dasatinib treatment effect on RBC deformability. **(A)** Illustration of the limited exponential growth function fit, used to characterize the shape probability curves presented on the control data. **(B)** Fitted growth rate for normocytes of control samples vs samples from ChAc patients. P-value calculated by Welch’s t-test (p = 0.009). **(C)** Fitted growth rate λ of the shape probability curves from ChAc patients as a function of treatment time with dasatinib for normocytes and **(D)** acanthocytes respectively. Dashed lines represent datapoints taken after treatment was stopped. Errorbars represent standard error of the fit, calculated from the covariance matrix. Gray region shows the respective value from healthy donors without treatment (Mean ± SD). **(E)** Percentage of acanthocytes found in each sample during shape analysis measurements for patients P1-P5. Dashed lines indicate datapoints taken after the respective treatment already stopped. **(F)** Fraction of healthy shapes (hs) and acanthocytes (ac) over cell velocity for untreated and *in-vitro* dasatinib treated RBCs from patient P3.

The shape analysis method is not suited to measure the deformability of WBCs, because the stresses acting the cells are too low and also the throughput of the method is not high enough to capture a sufficient number of cells.

### 2.3 Real-time fluorescence and deformability cytometry (RT-FDC)

The setup of real-time fluorescence and deformability cytometry (RT-FDC) is described in detail in Otto et al., 2015 (29) and Rosendahl et al., 2018 (30). In brief, flow is introduced into a microfluidic chip with the help of syringe pumps and cells get deformed by hydrodynamic stresses as they pass a narrow constriction, still larger than their own size. The chip is mounted on an inverted microscope (Axiovert 200M, ZEISS, Oberkochen, Germany) and images are obtained at the end of the constriction with a CMOS camera (Mikrotron, Unterschleissheim, Germany). Camera and syringe pump are controlled via the measuring software ShapeIn (Zellmechanik Dresden, Dresden, Germany). ShapeIn analyzes the recorded images in real-time and computes a contour and projected area for each cell. The deformation parameter used in RT-FDC is defined as 1-circularity and is calculated from the contour’s area and perimeter using: 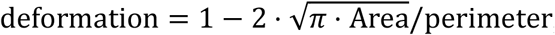, illustrated on the example of a deformed RBC in figure 1C. As introduced by Toepfner et al., 2018 (33), the data from RT-FDC experiments can be used to distinguish between different types of blood cells. These strategies were used here to selectively study the deformation and mechanics of RBCs, lymphocytes, and myelocytes from ChAc patients (see also SI text and fig. S5A,B).

Because the stresses required to deform RBCs are smaller than for leukocytes, both were measured under different conditions. For measurements on leukocytes, citrate blood was resuspended in a viscosity-adjusted measurement buffer (phosphate saline buffer without Mg^2+^ and Ca^2+^ (PBS−) containing 0.5% (w/v) methyl cellulose (4000 cPs, Alfa Aesar 036718.22, CAS# 9004-67-5); adjusted in HAAKE Falling Ball Viscometer type C (Thermo Fisher Scientific, Dreieich, Germany) using ball number 3 to a viscosity of 15 mPa s) at a ratio of 1:20. The increased viscosity of the buffer also increases the stresses acting on the cells inside the channel. Measurements were performed in channels with 20×20 µm cross-section at a flow rate of 0.08 µl/s. Example images of a deformed lymphocyte and myelocyte are shown in figure 1D and E, respectively. To decouple effects of the cell size on the deformation, the Young’s modulus was computed for leukocytes as described in Mokbel et al., 2017 (34). Since the derivation of Young’s moduli for this system is only valid for initially spherical cells (34,35), it cannot be calculated for RBCs.

RBC measurements were performed on an RT-FDC setup with a fluorescence module described in detail in Rosendahl et al., 2018 (30). Whole blood was resuspended in the same measurement buffer described above at a ratio of 1:200. The buffer was complemented with 2.5 µM syto13 nucleic acid stain (Thermo Fisher Scientific, Dreieich, Germany) to mark for the reticulocytes in the sample. Fluorescence was excited with a 60 mW, 488 nm laser at 8% power (OBIS 488-nm LS 60 mW, Coherent Deutschland) and the signal was measured with an avalanche photodiode (MiniSM10035; SensL Corporate, Cork, Ireland). Measurements were performed in channels with a 20×20 µm cross-section at a flow rate of 0.02 µl/s. The control data for the RT-FDC measurements on RBCs was not measured for this study but taken from 10 control samples for Rosendahl et al., 2018 (30) which were measured under the same conditions (data shown in figure S4E,F).

### 2.4 Data analysis and data availability

Data was analyzed and plotted using custom python scripts. A detailed documentation can be found here: https://gitlab.gwdg.de/freiche/changes-in-blood-cell-deformability-in-chorea-acanthocytosis-and-effects-of-treatment-with-dasatinib-or-lithium. A collection of the raw data files is on figshare: https://doi.org/10.6084/m9.figshare.c.5793482.

## 3 Results

### 3.1 Red blood cell deformability in Chorea-Acanthocytosis

To investigate differences in the deformability of red blood cells (RBCs) from chorea-acanthocytosis (ChAc) patients compared to that of healthy donors, we performed shape-analysis and real-time fluorescence and deformability cytometry (RT-FDC) measurements on the blood of 5 patients (P1-P5) and compared it to measurements of healthy controls. The results are depicted in figure 1A,B and 1F,G. Every velocity bin of 0.25 mm/s in figure 1A and B includes approx. 100 cells for each patient. The shape-probability curves of normocytes (fig. 1A) and acanthocytes (fig. 1B) show that RBCs from ChAc patients are less likely to deform in channel flow. It should be mentioned that also control samples can include acanthocyte-like cells but to a much lesser extent than in patient samples. The analysis for the control samples only included normocytes. Acanthocytes are clearly less likely to deform due to flow even for higher flow velocities.

Analysis of the exponential growth rate (see fig. 2A) of control and ChAc normocytes is depicted in figure 2B and showed that the growth rate for control normocytes was higher which indicated that the cells are were deformable. This is in line with findings from Rabe et al., 2021 (32) which reported similar results for ChAc normocytes. A comparison for acanthocytes was not possible because they were not present in sufficient numbers in control samples.

The RT-FDC measurements showed that the RBCs of ChAc patients had a slightly larger projected area than those from healthy donors (fig. 1F, effect size of +1.89 µm^2^, p=0.0036 computed with linear mixed effect models (36)). No significant difference between control and ChAc RBCs’ deformation could be detected (p=0.1382, fig. 1G).

### 3.2 White blood cell deformability in Chorea-Acanthocytosis

In RT-FDC measurements on whole blood it is possible to distinguish different cell types (see materials and methods, SI text and Töpfner et al., 2018 (33)). Thus, we investigated mechanical differences also between the leukocytes of ChAc patients and one healthy donor. We distinguished, mainly by size, between all lymphocytes and cells resulting from the myeloid lineage without RBCs (myelocytes), comprising all mono- and granulocytes. The myelocyte fraction mainly consisted of neutrophil granulocytes (>80%).

Since white blood cells are spherical at rest, we can employ a model that maps the projected area and the deformation to an apparent Young’s modulus (34), to see differences in stiffness before, during and after the treatments. As depicted in figure 1H,I, both lymphocytes and myelocytes of ChAc patients were significantly stiffer than their healthy counterparts before treatment onset (lymphocytes: +240 Pa, p=0.0186, myelocytes: +150 Pa, p=0.0070; effect sizes and p-values determined by linear mixed effects model analysis (36)). The projected area and deformation data used to compute the Young’s modulus is shown in figure S5C,D.

### 3.3 Dasatinib and lithium treatment effect on RBC deformability

#### 3.3.1 RBC shape analysis during dasatinib or lithium treatment

To study the effect of dasatinib or lithium on RBC deformability of ChAc patients, we monitored RBCs during treatment with shape analysis measurements and RT-FDC.

To compare shape-analysis data over the treatment time, a limited exponential growth was fitted to the data (see fig. 2A) and the fitted growth rate was used to detect changes in the deformability during the treatment time. The results for the dasatinib treatment are shown in figure 2C,D, those of the lithium in figure 3. Even though one might get the impression that the growth rate showed a slight increase with dasatinib treatment time for both normocytes and acanthocytes, the overall RBC deformability remained unaffected by the treatment. The results pooled for all RBCs are shown in figure S1 and the curves from all measurements are given in figure S2.

**Figure 3:**
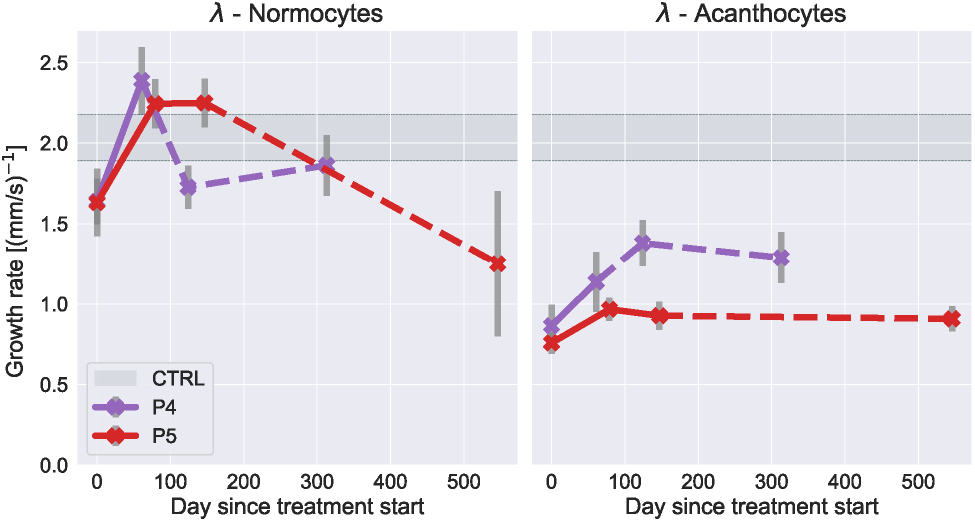
Lithium treatment effect on RBC deformability. Fitted growth rate λ of the shape probability curves from ChAc patients as a function of treatment time with lithium for normocytes and acanthocytes respectively. Dashed lines represent datapoints taken after treatment was stopped. Errorbars represent standard error of the fit, calculated from the covariance matrix. Gray region shows the respective value from healthy donors without treatment (Mean ± SD).

The acanthocyte count over the treatment is given in figure 2E and shows that the fraction of acanthocytes remained constant within a certain range of fluctuation without a systematic trend for both treatments, which was in line with the clinical data (19).

To check if dasatinib had a short-term effect on RBC deformability, RBCs from P3 were treated with dasatinib *in vitro* and the shapes in channel flow were analyzed (for details, see SI text). The results are shown in figure 2F. There was no significant difference in the deformability between control- and dasatinib-treated RBCs. This indicated that the observed effects on cell deformability were likely caused by long-term treatment effects, e.g., acting on erythropoiesis or indirect systemic effects.

For the lithium treatment, normocyte growth rate even increased above control values during the treatment and recovered to, or below, pre-treatment levels after the treatment was discontinued. Acanthocytes showed a slight increase in growth rate during lithium treatment. This indicated that the RBCs got softer during lithium treatment but RT-FDC results showed that the cells also got larger with the treatment (see fig. 4) which would make the cells more likely to deform in the channel if the overall mechanics are unchanged.

**Figure 4:**
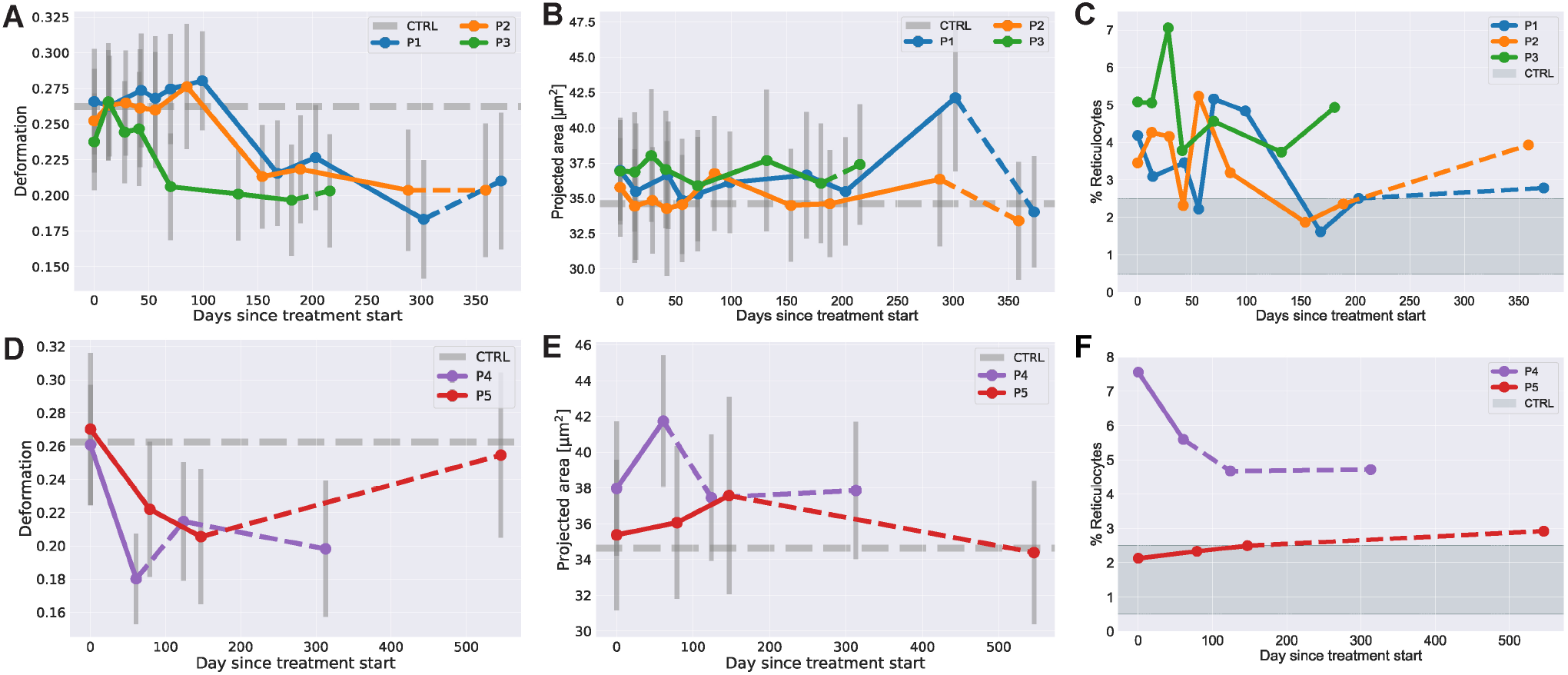
RBC deformation, size and reticulocyte count in RT-FDC during treatment with dasatinib or lithium. **(A-C)** RBC median deformation, projected area and reticulocyte fraction measured with RT-FDC before, during and after treatment with dasatinib. **(D-F)** RBC median deformation, projected area and reticulocyte fraction measured with RT-FDC before, during and after treatment with lithium. Dashed lines indicate time points after the treatment was stopped, dashed gray lines indicate control values without treatment. The red region in the reticulocyte plots indicates the range for healthy individuals reported in the literature of 0.5-2.5%.

#### 3.3.2 RBC deformation in RT-FDC during dasatinib or lithium treatment

RT-FDC results for the dasatinib treatment are shown in figure 4A-C and for the lithium treatment in figure 4D-F. It can be seen that during treatment with dasatinib, the RBC deformation parameter dropped for all patients between 50-100 days after treatment start, while the projected area remained mainly constant with the exception of some outliers.

For the lithium treatment, the RT-FDC results showed that the deformation parameter decreased with treatment time which was accompanied by an increase in cell size. After the treatment, the deformation and area values returned almost to the control values for P5 while the values for P4 did not fully recover.

The interpretation of deformation values for RBCs in RT-FDC cannot directly be linked to cell mechanics because they undergo a non-trivial shape transition from discocyte to tear-drop shape (28). A decrease of the deformation value does not necessarily mean that the cells got stiffer. To back this up, we artificially stiffened RBCs from healthy donors with glutaraldehyde (37,38) or diamide (39) (see SI text and fig. S3) and saw that an increasing concentration of the reagents led to an increase of the deformation when working at low concentrations. At high concentrations the cell shape got fixed which led to lower deformation values.

Since reticulocytes have been reported to have a higher projected area and lower deformation compared to the whole RBC population (30), we speculated that above mentioned results might be due to increased levels of reticulocytes in ChAc patients. The results for reticulocytes, labelled with syto13 nucleic acid stain, together with all RBCs are given in figure S4A-D. The patients’ reticulocytes indeed had higher projected area and lower deformation if compared to the whole RBC population which did not differ from healthy controls. Qualitatively, reticulocyte area and deformation changed over treatment time like the total RBC population. The fraction of reticulocytes in the blood was initially higher compared to controls and slightly decreased during the dasatinib treatment closer to values from healthy donors (fig. 4C). A reduction of the reticulocyte count with dasatinib treatment was also reported for VPS13a^-/-^ mice phenocopying human ChAc (18). During lithium treatment P4 showed a decrease during and after treatment, while P5 showed a slight increase but values were mostly in the range expected for healthy individuals (0.5-2.5%).

### 3.4 ChAc Leukocytes stiffened during treatment with dasatinib

The Young’s moduli of lymphocytes and myelocytes in dasatinib treatment are shown in figure 5A,B. The lithium data is shown in figure 5C,D. During treatment with dasatinib, we observed a further increase of the Young’s modulus for all three patients which also persisted for the lymphocytes shortly after the treatment stopped. For myelocytes, the median Young’s modulus decreased for patients P1 and P2 shortly after the treatment but kept increasing for P3.

**Figure 5:**
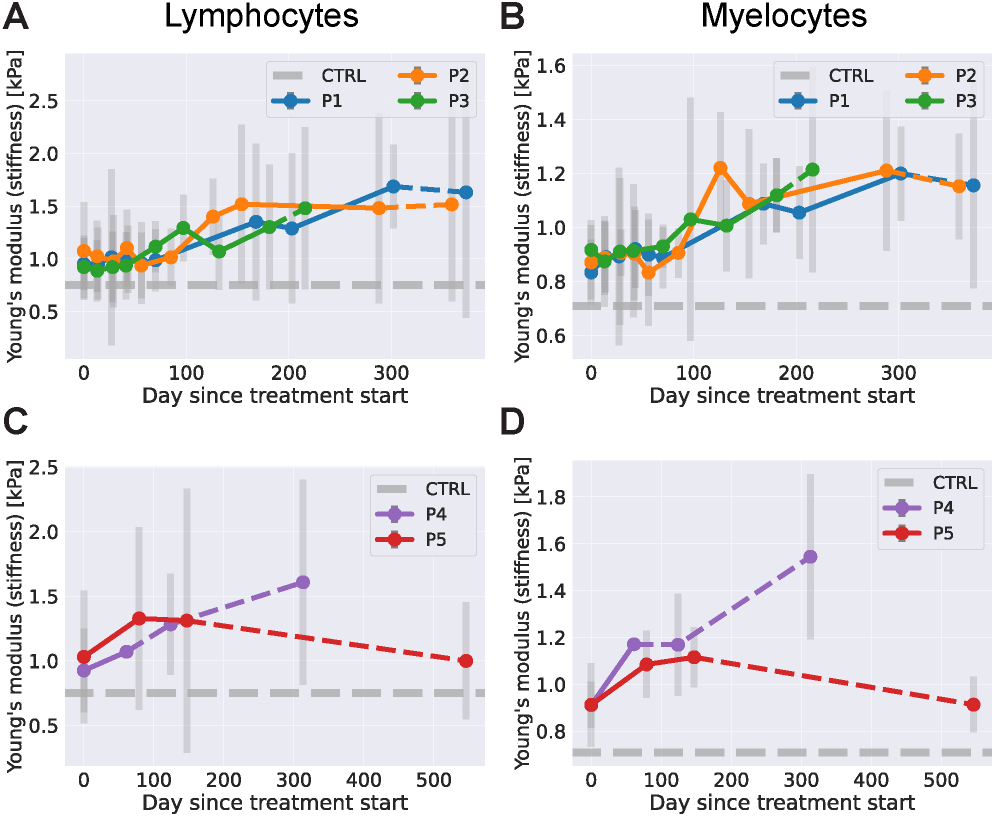
Leukocyte stiffness in RT-FDC during treatment with Dasatinib or lithium. **(A)** Median Young’s modulus of ChAc patients’ lymphocytes during dasatinib treatment. **(B)** Median Young’s modulus of ChAc patients’ myelocytes during dasatinib treatment. **(C)** Median Young’s modulus of ChAc patients’ lymphocytes during lithium treatment. **(D)** Median Young’s modulus of ChAc patients’ myelocytes during lithium treatment.

During treatment with lithium both patients showed an increase in lymphocyte and myelocyte stiffness. After the treatment was stopped, we observed a further increase for patient P4 and a drop back to the pre-treatment level for patient P5 approximately one year after the treatment stopped. The area and deformation data used to compute the Young’s modulus is given in figure S5E,F.

## 4 Discussion

Our study highlights that the deformability, not only of RBCs but of all blood cell types, was altered in ChAc patients. Shape analysis results showed a decreased deformability of ChAc RBCs compared to controls. The decreased deformability can be directly inferred from the reduced number of deformed cells at any cell velocity which was also reflected by a smaller growth rate. A reduced deformability of ChAc normocytes, deduced from cell shapes, has recently also been reported by Rabe et al., 2021 (32).

In RT-FDC, we did not observe a change in the deformation parameter but ChAc RBCs had a larger projected area. Because deformation and size are always correlated in RT-FDC (35), this is also an indicator for a change of RBC deformability, but the interpretation of this behavior is not straight forward. To see how the deformation parameter changes when RBC stiffness is controllably changed, we treated RBCs *in vitro* with glutaraldehyde or diamide. At low concentrations, we observed an increase of the deformation parameter which shows that higher deformation parameter can also occur for stiffer RBCs.

While we observed larger RBCs in ChAc patients, values for the mean corpuscular volume (MCV) reported in Peikert et al., 2021 (19) (see also Table S2) did not show an increased RBC volume for patients P1-P3. Sizes reported in RT-FDC measurements are based on the projected cross-sectional area of the deformed cells. Simulations of the flow fields inside the channel suggest that the stresses near the centerline can be approximated as axisymmetric (40). Therefore, it is fair to assume that the RBCs take up an axisymmetric shape as well and the projected area is directly correlated to cell size. In automated cell counters, the MCV is determined, e.g., by impedance or light scattering measurements which require a constant deformability of RBCs for a reliable result. Thus, MCV is not a direct measurement of single cell sizes. Here, we provide a direct observation of sizes based on single cell imaging.

Both, shape analysis and RT-FDC indicated a change of RBC deformability during treatment with dasatinib. Shape analysis experiments showed a slight increase of the growth rate with treatment time, which indicated an increased deformability of both normocytes and acanthocytes. In RT-FDC, we observed a sharp drop of the deformation parameter after 50-100 treatment days while the size fluctuated around a constant value. Again, for RBCs a direct link between the deformation parameter in RT-FDC and cell mechanics is not trivial and a decrease in deformation does not necessarily mean that the cells got stiffer (see fig. S3). Even though we do not have certainty of the direction of the change, we claim that RBC mechanics was altered during dasatinib treatment.

During lithium treatment, we also observed an increasing growth rate in the shape analysis which indicated increased RBC deformability. In RT-FDC, we saw a decrease of deformation accompanied by an increase in projected area. An increase in projected area and decrease in deformation hints at a volume increase and a reduced surface area-to-volume ratio, which leads to an increased sphericity or effective rounding of the cells. Rounder shapes lead to smaller deformation values, assuming that this effect dominates any stiffness changes of the cells. The increased cell size could also explain the increased deformability in the shape analysis experiments because larger cells experience higher stresses and are therefore more likely to deform. All in all, these findings indicate that lithium affects RBC properties in ChAc patients but because of the scarcity of the data here, one should be careful with the interpretation.

Data from Peikert et al., 2021 indicated that with dasatinib treatment, F-actin became more localized at the cortex (19). These modifications in the actin network likely contribute to changes in RBC deformability (41) that we observed here.

Treatment with dasatinib led to a decrease of the reticulocyte fraction in the blood of ChAc patients. The same effect was observed in a mouse model phenocopying human ChAc (18). Peikert et al., 2021 reported an abundance of the autophagy initiator Ulk1 in ChAc patients RBCs, which decreased with dasatinib treatment (19). Ulk1 is connected to autophagy in erythropoiesis and could influence the rate of reticulocyte production (42), providing a mechanistic link to the reticulocyte count.

While an impaired RBC deformability was described before for patients with other forms of neuroacanthocytosis (43) this is, to our knowledge, the first time that also patients’ leukocyte stiffness was studied. Lymphocytes and myelocytes were both stiffer in ChAc patients compared to cells from healthy donors. This is of interest as VPS13A, the gene which is known to have a loss-of-function mutation in ChAc, is especially expressed in human monocytes and B-, and helper T-cells (44). Since VPS13A is known to cause structural reorganizations of the cytoskeleton from studies on other cell types (17,20,45), it is likely that we observed these as altered mechanical properties in leukocytes. Another feature in ChAc blood is an increased expression of Lyn kinase, which is known to set the threshold for B-cell activation. Mechanical changes of lymphocytes after activation have been reported before (33,46). Another mechanism that could lead to altered leukocyte mechanics is an increase in interleukin-1 beta (IL-1b) that was observed in the brain of Vps13a knockout mice (18). IL-1b not only plays a key role in autoinflammation but is also known to cause re-organization of the actin cytoskeleton in different cell types (47,48).

Young’s moduli of both lymphocytes and myelocytes increased even further away from control values during treatment with dasatinib or lithium. A stiffening was previously also observed for leukemic cells during treatment with different chemotherapeutic agents for acute lymphoblastic leukemia (dexamethasone and daunorubicin), such as dasatinib (49). Dasatinib interferes with all the pathways described above and likely induces further mechanical changes in leukocytes. The inhibited activity of Lyn during dasatinib treatment could cause a change in lymphocyte activation levels leading to altered mechanics but the exact mechanisms for the observed leukocyte stiffening with dasatinib treatment remain rather speculative at this point.

Overall, our results showed that the mechanical phenotype of all blood cell types in ChAc patients were impaired but did not change towards regimes of healthy donors during the treatments. Clinical data in Peikert et al., 2021 (19) hints on slightly improved hematological features but these did not manifest in an improved neurological phenotype. This demonstrates that the techniques that were utilized here are suited to measure a treatment response.

In general, decreased blood cell deformability often leads to clinical symptoms due to decreased cell survival and secondary pathology (25–27,50). Especially, an impaired oxygen transport by altered blood cell deformability should be considered in neurological diseases (51). While clinical descriptions of ChAc patients focus on neurological symptoms, we highlight in this study that the effects on blood deformability should not be neglected and could contribute to the clinical manifestation (through the mechanisms stated above) and thus methods that are able to monitor this (as shown here) should be considered as read outs for clinical trials. Our results on the impaired mechanics of leucocytes especially highlights that the focus of ChAc research should not only be on RBCs but also include other blood cell types.

## Supporting information

Movie_S1_normocyte_undeformed_1

Movie_S2_normocyte_undeformed_2

Movie_S3_normocyte_deformed_1_slipper

Movie_S4_normocyte_deformed_2_parachute

Movie_S5_acanthocyte_undeformed_1

Movie_S6_acanthocyte_undeformed_2

Movie_S7_acanthocyte_deformed_1

Movie_S8_acanthocyte_deformed_2

## 6 Author Contributions

Conceptualization: F.R., M.K., K.P., G.B., A.H., and J.G.; Experiments–shape analysis: F.R. and A.R.P., Experiments–RT-FDC: F.R., P.R., M.H., and M.K., Experiments–in-vitro shape analysis: A.K.; Formal analysis: F.R., P.R., A.K., and M.K.; Supervision: L.K.,A.H., and J.G.; Writing– original draft, K.P. and F.R..; Writing–review and editing, F.R., M.K., K.P., G.B., L.K., A.H., and J.G.; All authors have read and agreed to the published version of the manuscript.

## 8 Acknowledgments

The authors thank all patients and their families for giving consent for publication of the data and the healthy control subjects for participation in this study. We are especially grateful to Glenn (†) and Ginger Irvine as the founders of the Advocacy for Neuroacanthocytosis Patients (www.naadvocacy.org) and to Susan Wagner and Joy Willard-Williford as representatives of the NA Advocacy USA (www.naadvocacyusa.org).

## 9 Supplementary Material

Supplementary Information, Figures, Table and Videos will be available online.

## 10 Data Availability statement

Raw data files and initial analysis scripts are available on figshare.com: https://doi.org/10.6084/m9.figshare.c.5793482. Analysis documentation and creation of figure plots is documented on gitlab: https://gitlab.gwdg.de/freiche/changes-in-blood-cell-deformability-in-chorea-acanthocytosis-and-effects-of-treatment-with-dasatinib-or-lithium.

