## Supplementary Information for "Changes in blood cell deformability in Chorea-Acanthocytosis and effects of treatment with dasatinib or lithium"

### 1 Supplementary Information

#### 1.1 RT-FDC gating strategies

Lymphocytes and myelocytes are distinguished by size like shown in figure S5A. In RT-FDC leucocyte measurements, red blood cells are not used in the analysis and filtered out in a first step by only allowing cells with an aspect ratio (x-size/y-size) smaller than 2. For ChAc patients, a lot of RBCs were found in the deformation-size region described for leucocytes. To exclude these events from the analysis, additionally all measurements were gated for brightness values within the contour, fitting for lymphocytes or myelocytes respectively depicted for one exemplary measurement in figure S5B (for a detailed explanation of the gating strategy, see Töpfner et al., 2018 (1)). Additionally, the contours are checked for convexity and an event is only considered in the WBC analysis if the area of the convex hull of the contour deviates by a maximum of 5% from the area of the raw contour. For RBC measurements a deviation by 10% was allowed and events were filtered in a size range of 20-50 $\mu m^2$.

#### 1.2 Shape Analysis of *in-vitro* Dasatinib treated RBCs

Blood samples were obtained via venipuncture and collecting the blood in vacutainers containing the anticoagulant heparin. An amount of approximately 100 µl was suspended in 1ml phosphate buffered saline (PBS) and centrifuged at 1500 g for 5 min. After this centrifugation, the supernatant was discarded and the residual pellet of RBCs is resuspended in PBS. This process is repeated three times in total, leaving a dense pellet of RBCs devoid of blood plasma and other cellular constituents. To obtain the final blood solution used in the microfluidic experiments, a volume of 10 µl of these RBCs was filled to 1 ml total volume with a solution of PBS and 1 mg/ml bovine serum albumin (BSA), yielding a hematocrit of 1%.

For the Dasatinib measurements, a stock solution of 0.5 mg Dasatinib per 1 ml dimethyl sulfoxide (DMSO), yielding a concentration of ca. 1 µmol/l, was created. From this stock solution, an additional amount of 10 µl was then added to the formerly described final blood solution. In addition, the RBCs were incubated in 10 µl of the previously described stock solution, filled to 1ml total volume with PBS, at room temperature for at least 3 h prior the actual measurement.

By applying a set of discrete pressure drops in a regime up to 700 mbar to a connected Eppendorf tube containing the final blood solution, we advected this solution through microfluidic channels with a cross-section of (w x h) 12 µm x 10 µm, and a length of ca. 4 cm.

The flowing RBCs were recorded with a brightfield microscope equipped with an oil-immersion objective (Nikon Plan Apo 60x) at a distance 10 mm away from the channel entrance to neglect transient cell shapes (cf. references (2,3) for more information). With the aid of a custom-tailored particle tracking algorithm, we extract cropped images of single flowing RBCs together with their individual velocities and centroid positions.

Cells flowing due to a distinct pressure drop are grouped and from this set the mean velocity was obtained. Analogously, cells associated to one distinct pressure drop were manually categorized into a binary scheme of healthy shapes (hs) and acanthocytes (ac).

The results are shown in main text figure 2F. It can be seen that the number of acanthocytes decreases with the flow velocity. This is because the typical morphologic acanthocyte features can vanish with cell deformation. There was no significant difference between the control and Dasatinib-treated sample.

### 1.3 RBC treatment with glutaraldehyde and diamide

To see how the deformation of RBCs changes when artificially altering the cells' material properties, we incubated them with different concentrations of glutaraldehyde (0.0001-0.005 v%; Sigma-Aldrich, Cat# G6257) or diamide (0.1-10 mM, Sigma-Aldrich, Cat# D3648) for 20 min in the measurement buffer before measuring. While glutaraldehyde crosslinks proteins throughout the whole cell, diamide specifically targets the spectrin network at the RBC cytoskeleton. The results for the measured projected areas and deformations are given in figure S3.

For glutaraldehyde, cells decrease in size for concentrations up to 0.001 v% with remaining or slightly increasing deformation values. At the concentration of 0.0025 v% most cells are fixed in shape, which leads to two populations for the measured size (dependent on the direction of the cell when the image is taken) and also for the deformation, while deformation values clearly drop at these concentrations.

RBCs show a decrease in measured area with an increasing diamide concentration. For all concentrations shown here, we observe the characteristic tear drop shape (see main text fig. 1C) for RBCs in RT-FDC. Up to a concentration of 0.5 mM we see a deformation increase compared to control cells. At 1 mM and over, deformation values are smaller compared to control cells.

This highlights that a stiffening of RBCs not necessarily leads to a decrease of the deformation value in RT-FDC and even the opposite can be observed.

### 1.4 Data analysis

RT-FDC data was analyzed with the tools dclab (4) and ShapeOut2 (5).

Shape analysis experiments were analyzed using custom built cell tracking software in python 2.7 (6) and further analysis was performed in python 3.7 (7) mainly using the packages numpy (8), matplotlib (9), pandas (10) and seaborn (11).

6. Van Rossum G, Drake Jr FL. Python reference manual. Centrum voor Wiskunde en Informatica Amsterdam; (1995.).

7. Van Rossum, Guido and Drake FL. Python 3 Reference Manual. CreateSpace; (2009.).

8. Harris CR, Millman KJ, van der Walt SJ, Gommers R, Virtanen P, Cournapeau D, et al. Array programming with NumPy [Internet]. Vol. 585, Nature. Nature Research; (2020.) p. 357–62. doi: 10.1038/s41586-020-2649-2

9. Hunter JD. Matplotlib: A 2D graphics environment. Comput Sci Eng. (2007) 9(3):90–5.

10. McKinney W. Data Structures for Statistical Computing in Python. In: Proceedings of the 9th Python in Science Conference. SciPy; (2010.) p. 56–61.

11. Waskom M. seaborn: statistical data visualization. J Open Source Softw. (2021) 6(60):3021. doi: 10.21105/joss.03021

### 2 Supplementary Movies

Movies S1 & S2: Examples of undeformed normocytes flowing through the channel

Movies S3 & S4: Examples of deformed normocytes

Movies S5 & S6: Examples of undeformed acanthocytes

Movies S7 & S8: Examples of deformed acanthocytes

### 3 Supplementary Figures and Tables

#### 3.1 Supplementary Figures

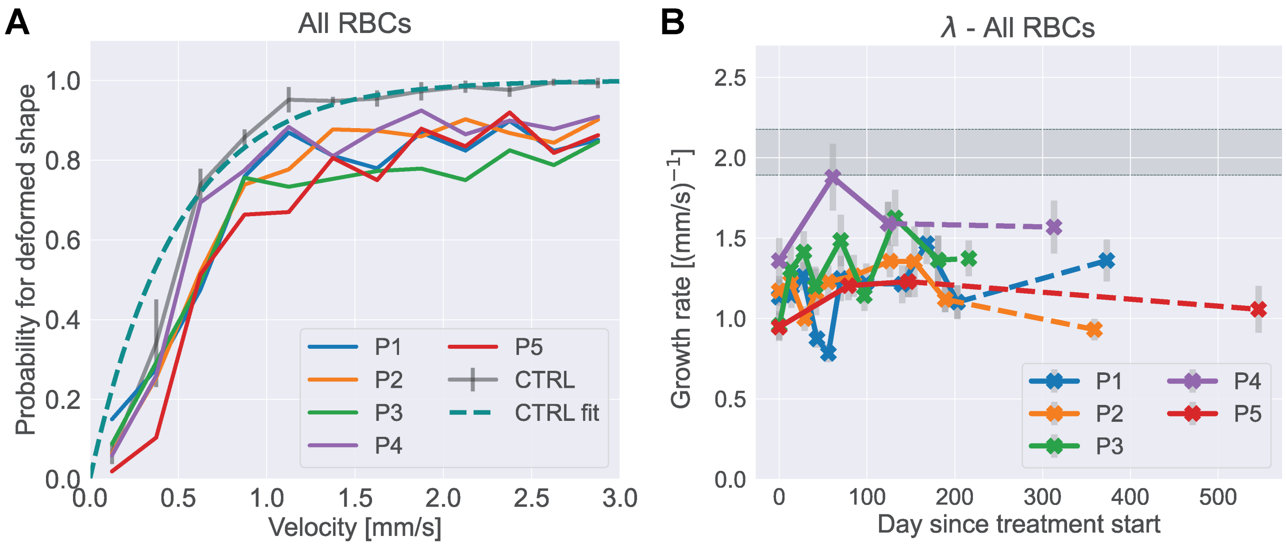


**Supplementary Figure 1. Shape Analysis – additional plots. (A)** Probability for deformed cells over flow velocity including all RBCs (normocytes and acanthocytes) from ChAc patients before treatment. **(B)** Fitted growth rate $\lambda$~~plateau probability $A$~~ for all RBCs during Dasatinib or Lithium treatment. Dashed lines represent datapoints taken after treatment was stopped. Errorbars represent standard error of the fit, calculated from the covariance matrix. Gray region shows the respective value from healthy donors without treatment (Mean ± SD). ~~**(C)** Fitted transition velocity $v_0$ for all RBCs during Dasatinib or Lithium treatment. **(D)** Fitted growth rate $k$ for all RBCs during Dasatinib or Lithium treatment. Dashed lines indicate datapoints taken after the respective treatment already stopped. Dashed gray lines indicate control values.~~

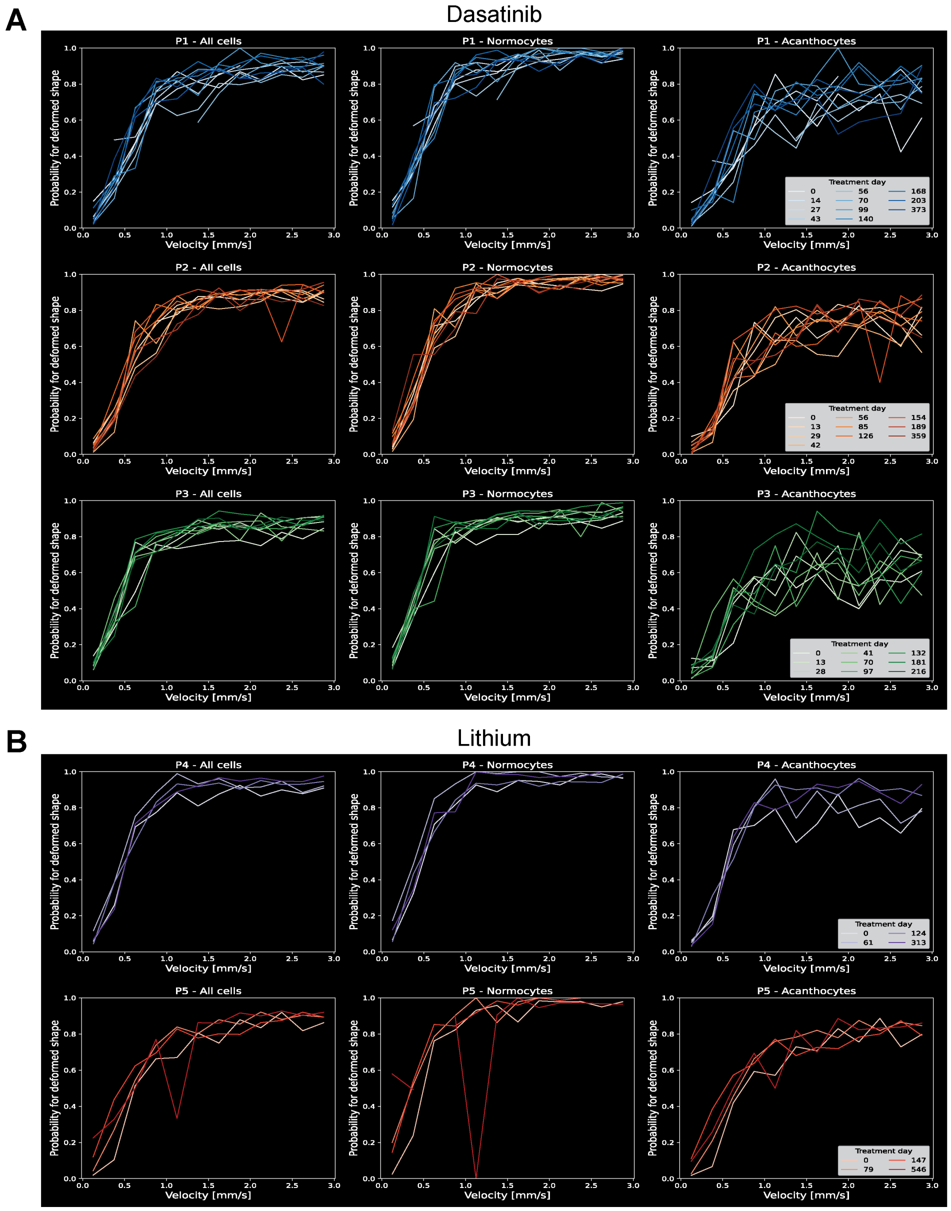


**Supplementary Figure 2.** Shape probability curves for all patients at every day of treatment with **(A)** Dasatinib or **(B)** Lithium (day 0 is before treatment start).

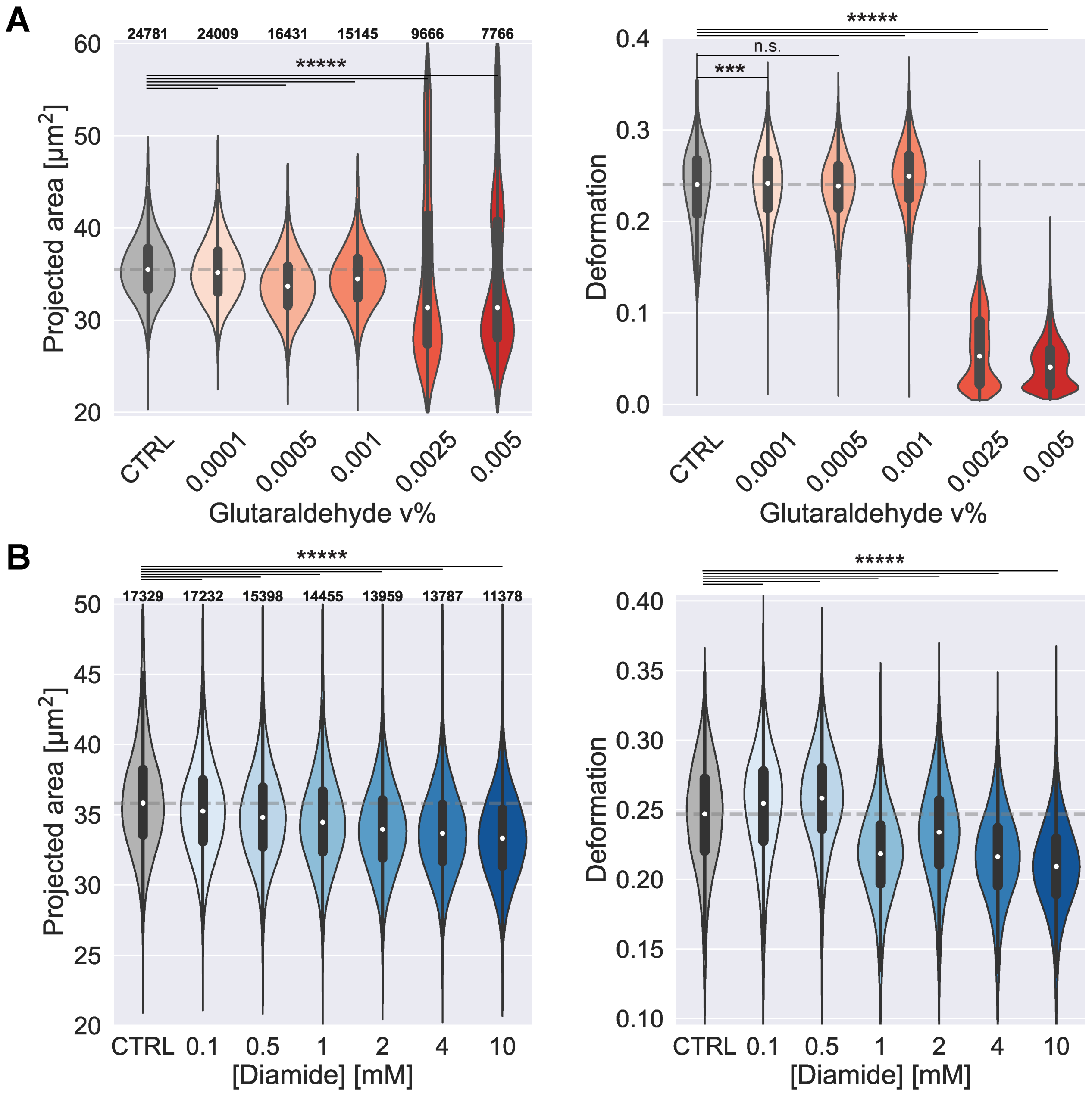


**Supplementary Figure 3. Effects of interference with RBC cytoskeleton in RT-FDC. (A)** Projected area and deformation of RBCs treated with different concentrations of glutaraldehyde. P-values calculated with Mann-Whitney U-Test. Projected area: all p-values < $10^{-5}$, deformation: CTRL – 0.0001%: p=0.0004; – 0.0005%: p=0.07; rest p< $10^{-5}$. Numbers on top of the plots indicate the number of observations per violin. **(B)** Projected area and deformation of RBCs treated with different concentrations of diamide. Dashed gray lines indicate median control values. All p-values < $10^{-5}$

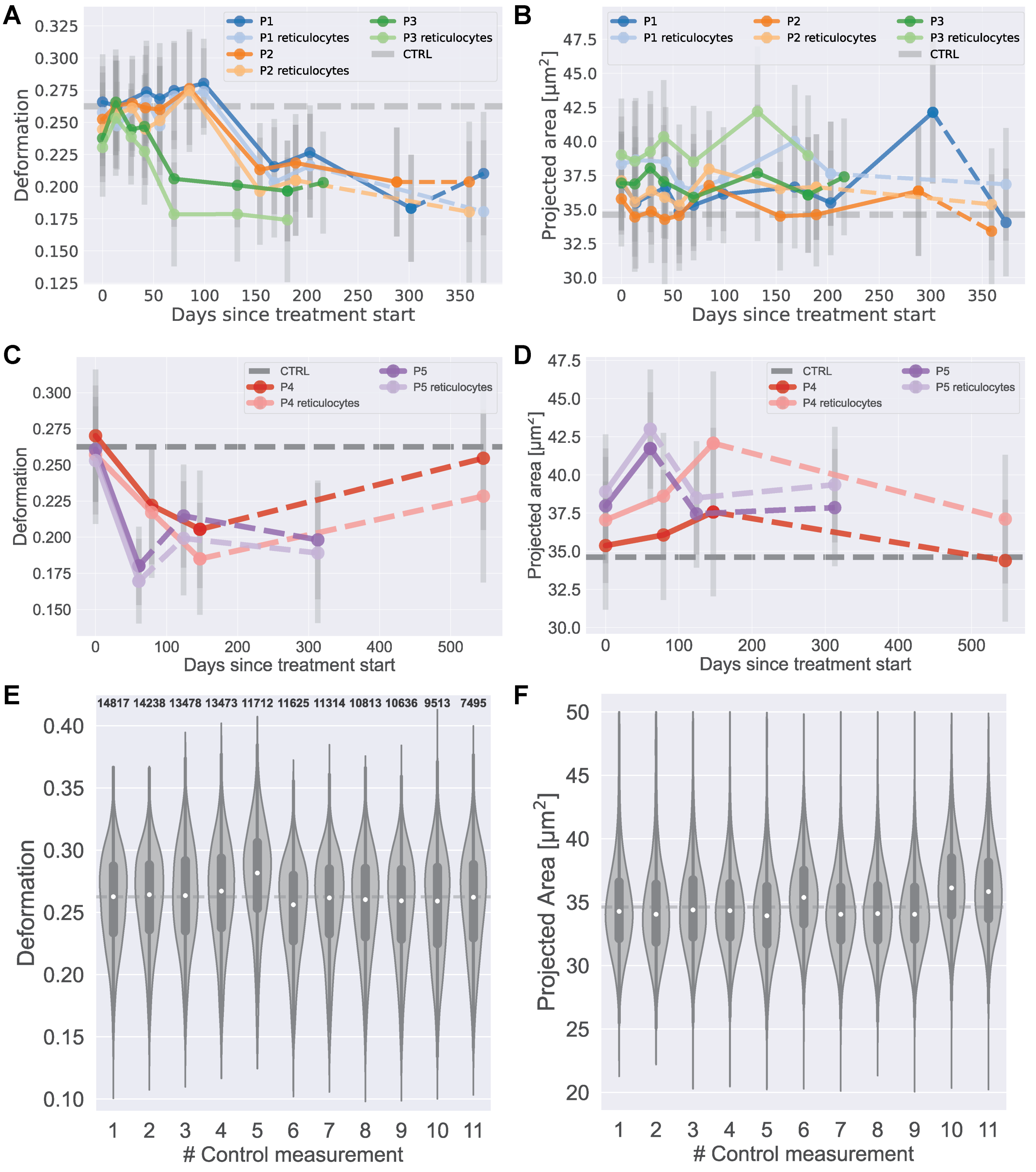


**Supplementary Figure 4.: RT-FDC on RBCs. (A, B)** Deformation and Projected area of all RBCs and syto13-positive reticulocytes during treatment with dasatinib. **(C, D)** Deformation and Projected area of all RBCs and syto13-positive reticulocytes during treatment with lithium. **(E, F)** Deformation and projected area for RBCs in control measurements from healthy donors. Numbers on top of the plots indicate the number of observations per violin.

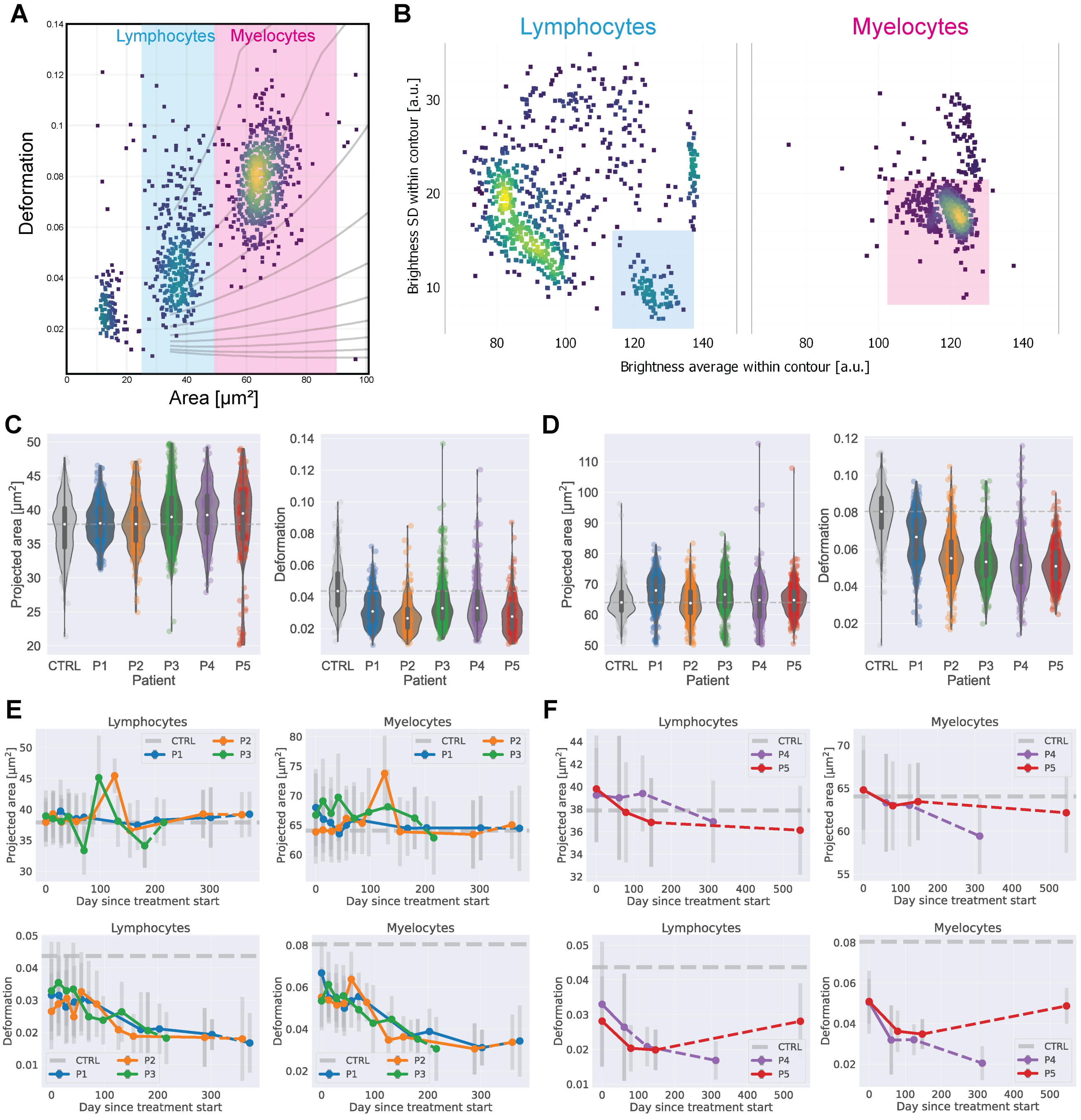


**Supplementary Figure 5. RT-FDC on white blood cells (A)** Gating for lymphocytes or myelocytes by size shown on a sample from a healthy donor. The population of cells smaller than 20 µm$^2$ is due to thrombocytes. **(B)** Excluding RBCs falling within the lymphocyte or myelocyte fraction by the standard deviation and average of brightness values within the contour. Cells not falling in the marked region are mainly single or aggregated RBCs. Shown here is a representative sample from patient P1.

**(C)** Projected area and deformation of lymphocytes of ChAc patients vs controls without treatment. **(D)** Projected area and deformation of myelocytes of ChAc patients vs controls without treatment. **(~~C~~E)** Projected area and deformation ~~Deformation and projected area~~ of lymphocytes and myelocytes during dasatinib treatment. **(~~D~~F)** Projected area and deformation ~~Deformation and projected area~~ during lithium treatment for lymphocytes and myelocytes.

**Supplementary Table 1: Clinical parameters of patients.**

| | | Sex/<br>Age (at d0) | Clinical characteristics | Time on treatment |
|---|---|---|---|---|
| Dasatinib | **P1** | ♂<br>27 years | Epilepsy<br>Mild chorea<br>Cognitive decline | **353 days** |
| | **P2** | ♂<br>32 years | Epilepsy<br>Mild chorea<br>Mild cognitive decline<br>Muscle atrophy | **341 days** |
| | **P3** | ♀<br>49 years | Parkinsonism<br>Epilepsy<br>Cognitive decline | **181 days** |
| Lithium | **P4** | ♂<br>52 years | Parkinsonism<br>Dystonia<br>Mild depression | **147 days** |
| | **P5** | ♂<br>53 years | Parkinsonism<br>Dystonia<br>Cognitive decline<br>Epilepsy | **71 days** |

### Supplementary Table 2: Complete blood count, RBC indices and hemolytic parameters.

| | normal range | unit | P1 | | | | | | | | | | | P2 | | | | | | | | | |
|---|---|---|---|---|---|---|---|---|---|---|---|---|---|---|---|---|---|---|---|---|---|---|---|
| Time after treatment start | | days | 0 | 14 | 27 | 43 | 56 | 70 | 99 | 168 | 203 | 302 | 373 | 0 | 13 | 29 | 42 | 56 | 85 | 154 | 189 | 288 | 359 |
| On treatment | | | No | Yes | Yes | Yes | Yes | Yes | Yes | Yes | Yes | Yes | No | No | Yes | Yes | Yes | Yes | Yes | Yes | Yes | Yes | No |
| Hemoglobin | 8,60 - 12,10 | mmol/L | 9.20 | 8.80 | 9.20 | 9.00 | 8.80 | 9.00 | 9.00 | 9.60 | 10.00 | 9.80 | 8.70 | 9.30 | 9.00 | 9.20 | 8.70 | 8.70 | 9.00 | 8.60 | 9.10 | 9.60 | 8.80 |
| Hematocrit | 0,400 - 0,540 | . | 0.42 | 0.40 | 0.40 | 0.41 | 0.40 | 0.41 | 0.40 | 0.44 | 0.45 | 0.44 | 0.40 | 0.43 | 0.42 | 0.41 | 0.40 | 0.40 | 0.42 | 0.40 | 0.41 | 0.43 | 0.39 - |
| White cell count | 3,8 - 9,8 | GPt/L | 5.80 | 8.45 | 6.81 | 6.32 | 6.61 | 7.33 | 5.72 | 7.16 | 5.45 | 7.19 | 5.56 | 7.61 | 8.23 | 6.45 | 7.63 | 7.44 | 7.46 | 4.76 | 6.18 | 6.92 | 5.52 |
| Plateled count | 150 - 400 | GPt/L | 172 | 159 | 139 - | 167 | 140 - | 112 - | 133 - | 189 | 200 | 158 | 160 | 242 | 147 - | 211 | 177 | 217 | 150 | 217 | 220 | 163 | 182 |
| Red cell count | 4,60 - 6,20 | TPt/L | 4.90 | 4.66 | 4.63 | 4.74 | 4.63 | 4.69 | 4.51 - | 4.91 | 5.05 | 4.93 | 4.49 - | 5.11 | 4.95 | 4.90 | 4.69 | 4.66 | 4.85 | 4.64 | 4.76 | 5.01 | 4.61 |
| Mean corpuscular hemoglobin (MCH) | 1,70 - 2,10 | fmol | 1.88 | 1.89 | 1.99 | 1.90 | 1.90 | 1.92 | 2.00 | 1.96 | 1.98 | 1.99 | 1.94 | 1.82 | 1.82 | 1.88 | 1.86 | 1.87 | 1.86 | 1.85 | 1.91 | 1.92 | 1.91 |
| Mean corpuscular hemoglobin concentration (MCHC) | 19,0 - 22,0 | mmol/L | 21.7 | 21.8 | 22.9 + | 21.7 | 21.7 | 21.8 | 22.2 + | 21.9 | 22.0 | 22.4 + | 21.6 | 21.5 | 21.3 | 22.3 + | 21.8 | 21.9 | 21.5 | 21.6 | 22.2 + | 22.5 + | 22.6 + |
| Mean corpuscular volume (MCV) | 80 - 96 | fl | 86 | 87 | 87 | 87 | 88 | 88 | 90 | 89 | 90 | 89 | 90 | 84 | 85 | 84 | 85 | 85 | 86 | 86 | 86 | 85 | 85 |
| Neutrophils (rel.) | 36,0 - 77,0 | % | 53.7 | 52.5 | 46.6 | 50.3 | 53.6 | 45.4 | 49.6 | 62.8 | 55.0 | 57.7 | 56.3 | 66.8 | 52.8 | 54.8 | 52.7 | 53.1 | 50.1 | 51.3 | 57.6 | 54.4 | 55.7 |
| Lymphocytes (rel.) | 20,0 - 49,0 | % | 36.8 | 38.1 | 43.6 | 38.9 | 37.2 | 44.7 | 40.0 | 32.7 | 33.4 | 32.1 | 31.7 | 22.1 | 36.0 | 31.5 | 33.9 | 35.3 | 36.1 | 35.5 | 30.6 | 34.5 | 33.5 |
| Monocytes (rel.) | 0,0 - 9,0 | % | 6.6 | 7.2 | 7.2 | 8.9 | 6.7 | 7.2 | 8.2 | 3.2 | 8.1 | 6.0 | 8.1 | 8.0 | 8.4 | 9.6 + | 9.8 + | 7.9 | 9.5 + | 9.2 + | 9.4 + | 8.7 | 8.5 |
| Eosinophils (rel.) | 0,0 - 5,0 | % | 2.2 | 1.8 | 2.3 | 1.6 | 2.0 | 2.2 | 1.9 | 0.7 | 2.9 | 3.6 | 3.2 | 2.6 | 1.9 | 3.3 | 3.1 | 3.0 | 3.6 | 3.2 | 1.9 | 1.4 | 1.4 |
| Basophils (rel.) | 0,0 - 1,0 | % | 0.7 | 0.4 | 0.3 | 0.3 | 0.5 | 0.5 | 0.3 | 0.6 | 0.6 | 0.6 | 0.7 | 0.5 | 0.9 | 0.8 | 0.5 | 0.7 | 0.7 | 0.8 | 0.5 | 1.0 | 0.9 |
| Neutrophils (abs.) | 1,80 - 7,55 | GPt/L | 3.14 | 4.44 | 3.17 | 3.18 | 3.55 | 3.32 | 2.83 | 4.50 | 3.00 | 4.15 | 3.13 | 5.08 | 4.35 | 3.54 | 4.01 | 3.95 | 3.74 | 2.44 | 3.56 | 3.76 | 3.07 |
| Lymphocytes (abs.) | 1,50 - 4,00 | GPt/L | 2.15 | 3.22 | 2.97 | 2.46 | 2.46 | 3.28 | 2.29 | 2.34 | 1.82 | 2.31 | 1.76 | 1.68 | 2.96 | 2.03 | 2.59 | 2.63 | 2.69 | 1.69 | 1.89 | 2.39 | 1.85 |
| Monocytes (abs.) | 0,20 - 1,00 | GPt/L | 0.36 | 0.61 | 0.49 | 0.56 | 0.44 | 0.53 | 0.47 | 0.23 | 0.44 | 0.43 | 0.45 | 0.61 | 0.69 | 0.62 | 0.75 | 0.59 | 0.71 | 0.44 | 0.58 | 0.60 | 0.47 |
| Eosinophils (abs.) | 0,00 - 0,49 | GPt/L | 0.12 | 0.15 | 0.16 | 0.10 | 0.13 | 0.16 | 0.11 | 0.05 | 0.16 | 0.26 | 0.18 | 0.20 | 0.16 | 0.21 | 0.24 | 0.22 | 0.27 | 0.15 | 0.12 | 0.10 | 0.08 |
| Basophils (abs.) | 0,00 - 0,20 | GPt/L | 0.03 | 0.03 | 0.02 | 0.02 | 0.03 | 0.04 | 0.02 | 0.04 | 0.03 | 0.04 | 0.04 | 0.04 | 0.07 | 0.05 | 0.04 | 0.05 | 0.05 | 0.04 | 0.03 | 0.07 | 0.05 |
| Bilirubin (total, serum) | < 21,0 | µmol/L | 5.2 | 4.9 | 5.4 | 6.5 | 6.8 | 4.7 | 4.4 | 5.0 | 6.0 | 4.1 | 3.4 | 5.9 | 5.4 | 4.5 | 5.7 | 4.8 | 4.6 | 4.1 | 4.7 | 6.2 | 5.3 |
| Haptoglobin (serum) | 0,30 - 2,00 | g/L | 0.32 | 0.68 | 0.23 - | 0.36 | 0.27 - | 0.44 | 0.32 | 0.57 | 0.54 | 0.51 | 0.11 - | 0.71 | 0.69 | 0.66 | 0.66 | 0.72 | 0.58 | 0.71 | 0.66 | 0.75 | 0.43 |
| Ferritin (serum) | 30,0 - 400,0 | µg/L | 73.9 | 100.3 | 73.6 | 66.8 | 75.6 | 59.8 | 45.6 | 46.6 | 28.0 - | 30.0 | 25.3 - | 145.7 | 132.4 | 135.9 | 116.2 | 118.2 | 105.5 | 95.3 | 70.3 | 81.3 | 61.3 |

| | normal range | unit | P3 | | | | | | | | P4 | | | | P5 | | | |
|---|---|---|---|---|---|---|---|---|---|---|---|---|---|---|---|---|---|---|
| Time after treatment start | | days | 0 | 13 | 28 | 41 | 70 | 132 | 181 | 216 | 0 | 61 | 124 | 313 | 0 | 79 | 147 | 546 |
| On treatment | | | No | Yes | Yes | Yes | Yes | Yes | Yes | No | No | Yes | No | No | No | Yes | Yes | No |
| Hemoglobin | 8,60 - 12,10 | mmol/L | 9.30 | 8.30 | 8.20 | 7.90 | 8.30 | 9.00 | 8.60 | 9.10 | 9.40 | 9.40 | 10.20 | 9.50 | 9.80 | 9.20 | 8.70 | 9.30 |
| Hematocrit | 0,400 - 0,540 | . | 0.43 | 0.39 | 0.38 | 0.36 - | 0.38 | 0.43 | 0.40 | 0.40 | 0.43 | 0.45 | 0.46 | 0.41 | 0.45 | 0.43 | 0.40 | 0.41 |
| White cell count | 3,8 - 9,8 | GPt/L | 5.02 | 5.31 | 4.96 | 5.35 | 5.08 | 6.22 | 4.52 | 4.60 | 4.61 | 6.32 | 4.04 | 4.84 | 7.13 | 8.99 | 7.30 | 7.63 |
| Plateled count | 150 - 400 | GPt/L | 137 - | 153 | 141 - | 132 - | 139 - | 156 | 158 | 125 - | 181 | 235 | 196 | 276 | 144 - | 129 - | 133 - | 133 - |
| Red cell count | 4,60 - 6,20 | TPt/L | 4.56 | 4.07 - | 3.94 - | 3.68 - | 3.80 - | 4.32 | 4.11 - | 4.31 | 4.54 - | 4.56 - | 4.92 | 4.58 - | 5.09 | 4.61 | 4.45 - | 4.66 |
| Mean corpuscular hemoglobin (MCH) | 1,70 - 2,10 | fmol | 2.04 | 2.04 | 2.08 | 2.15 + | 2.18 + | 2.08 | 2.09 | 2.11 | 2.07 | 2.06 | 2.07 | 2.07 | 1.92 | 2.00 | 1.96 | 2.00 |
| Mean corpuscular hemoglobin concentration (MCHC) | 19,0 - 22,0 | mmol/L | 21.8 | 21.2 | 21.4 | 21.8 | 21.8 | 21.1 | 21.5 | 22.7 | 22.0 | 21.1 | 22.4 + | 23.0 + | 21.8 | 21.4 | 21.9 | 22.9 + |
| Mean corpuscular volume (MCV) | 80 - 96 | fl | 93 | 96 | 97 + | 99 + | 100 + | 99 + | 97 + | 93 | 94 | 98 + | 92 | 90 | 88 | 93 | 89 | 87 |
| Neutrophils (rel.) | 36,0 - 77,0 | % | 42.0 | 35.1 - | 32.3 - | 33.5 - | 35.0 - | 29.0 - | 33.2 - | 34.2 - | 69.0 | 70.7 | 68.3 | 67.0 | 62.4 | 71.7 | 65.0 | 61.9 |
| Lymphocytes (rel.) | 20,0 - 49,0 | % | 45.2 | 51.6 + | 53.8 + | 52.5 + | 50.8 + | 56.4 + | 49.6 + | 50.9 + | 19.5 - | 16.5 - | 20.3 | 21.9 | 23.8 | 16.8 - | 23.8 | 22.8 |
| Monocytes (rel.) | 0,0 - 9,0 | % | 11.2 + | 10.5 + | 11.3 + | 12.1 + | 11.8 + | 12.4 + | 14.8 + | 13.0 + | 6.3 | 6.8 | 6.9 | 7.0 | 7.2 | 5.2 | 6.2 | 8.8 |
| Eosinophils (rel.) | 0,0 - 5,0 | % | 1.0 | 2.4 | 2.2 | 1.5 | 2.0 | 1.9 | 2.0 | 1.5 | 3.9 | 5.1 + | 3.0 | 2.9 | 5.8 + | 5.7 + | 4.2 | 5.6 + |
| Basophils (rel.) | 0,0 - 1,0 | % | 0.6 | 0.4 | 0.4 | 0.4 | 0.4 | 0.3 | 0.4 | 0.4 | 1.3 + | 0.9 | 1.5 + | 1.2 + | 0.8 | 0.6 | 0.8 | 0.9 |
| Neutrophils (abs.) | 1,80 - 7,55 | GPt/L | 2.11 | 1.86 | 1.60 - | 1.79 - | 1.78 - | 1.80 | 1.50 - | 1.57 - | 3.18 | 4.47 | 2.76 | 3.24 | 4.45 | 6.45 | 4.74 | 4.72 |
| Lymphocytes (abs.) | 1,50 - 4,00 | GPt/L | 2.27 | 2.74 | 2.67 | 2.81 | 2.58 | 3.51 | 2.24 | 2.34 | 0.90 - | 1.04 - | 0.82 - | 1.06 - | 1.70 | 1.51 | 1.74 | 1.74 |
| Monocytes (abs.) | 0,20 - 1,00 | GPt/L | 0.56 | 0.56 | 0.56 | 0.65 | 0.60 | 0.77 | 0.67 | 0.60 | 0.29 | 0.43 | 0.28 | 0.34 | 0.51 | 0.47 | 0.45 | 0.67 |
| Eosinophils (abs.) | 0,00 - 0,49 | GPt/L | 0.05 | 0.13 | 0.11 | 0.08 | 0.10 | 0.12 | 0.09 | 0.07 | 0.18 | 0.32 | 0.12 | 0.14 | 0.41 | 0.51 + | 0.31 | 0.43 |
| Basophils (abs.) | 0,00 - 0,20 | GPt/L | 0.03 | 0.02 | 0.02 | 0.02 | 0.02 | 0.02 | 0.02 | 0.02 | 0.06 | 0.06 | 0.06 | 0.06 | 0.06 | 0.05 | 0.06 | 0.07 |
| Bilirubin (total, serum) | < 21,0 | µmol/L | 7.7 | 5.9 | 6.3 | 5.3 | 5.1 | 5.3 | 5.6 | 5.8 | 12.1 | 11.3 | 12.2 | 17.9 | 9.0 | 8.5 | 10.5 | 12.5 |
| Haptoglobin (serum) | 0,30 - 2,00 | g/L | < 0.10 - | < 0.10 - | < 0.10 - | < 0.10 - | < 0.10 - | < 0.10 - | < 0.10 - | < 0.10 - | 0.13 - | 0.53 | 0.17 - | 0.44 | 0.50 | 0.97 | 0.49 | 0.28 - |
| Ferritin (serum) | 30,0 - 400,0 | µg/L | 150.3 | 145.2 | 126.6 | 116.5 | 110.7 | 123.4 | 107.7 | 74.0 | 49.0 | 65.8 | 39.0 | 75.4 | 16.8 - | 39.7 | 34.9 | 34.3 |
